# Identifying B-cell epitopes using AlphaFold2 predicted structures and pretrained language model

**DOI:** 10.1101/2022.12.06.519221

**Authors:** Yuansong Zeng, Zhuoyi Wei, Qianmu Yuan, Sheng Chen, Weijiang Yu, Yutong Lu, Jianzhao Gao, Yuedong Yang

## Abstract

**Motivation:** Identifying the B-cell epitopes is an essential step for guiding rational vaccine development and immunotherapies. Due to experimental approaches being expensive and time-consuming, many computational methods have been designed to assist B-cell epitope prediction. However, existing sequence-based methods have limited performance since they only use contextual features of the sequential neighbors while neglecting structural information.

**Results:** Based on the recent breakthrough of AlphaFold2 in protein structure prediction, we propose GraphBepi, a novel graph-based model for accurate B-cell epitope prediction. GraphBepi first generates the effective information sequence representations and protein structures from antigen sequences through the pretrained language model and AlphaFold2, respectively. GraphBepi then applies the edge-enhanced deep graph neural network (EGNN) to capture the spatial information from predicted protein structures and leverages the bidirectional long short-term memory neural networks (BiLSTM) to capture long-range dependencies from sequences. The low-dimensional representation learned by EGNN and BiLSTM is then combined to predict B-cell epitopes through a multilayer perceptron. Through comprehensive tests on the curated epitope dataset, GraphBepi was shown to outperform the state-of-the-art methods by more than 5.5% and 44.0% in terms of AUC and AUPR, respectively. We also provide the GraphBepi web server that is freely available at https://biomed.nscc-gz.cn/apps/GraphBepi.

**Availability:** The datasets, pre-computed features, source codes, and the pretrained model of GraphBepi are available at https://github.com/biomed-AI/GraphBepi.

## 1 Introduction

B-cells are a crucial element of the immune system, which could provide immunological protection against harmful molecules or infectious pathogens by producing antibodies that bind with antigens (Martin, et al., 2016). The specific region of an antigen binding to an antibody is known as antigenic determinant or an epitope(Paul, 2012). The category of B-cell epitopes (BCEs) is broadly classified into two groups: linear epitopes and conformational epitopes (Alghamdi, et al., 2022). Linear epitopes include continuous amino acid residues, whereas conformational epitopes are shaped by a three-dimensional conformation that folds the protein to bind through the interaction of discontinuous amino acid residues. Previous studies show that more than 90% of BCEs are discontinuous/conformational and 10% are linear epitopes (Barlow, et al., 1986).

Reliable tools for the identification of B-cell epitopes are important in many biotechnological and clinical applications, such as therapeutic antibody development and vaccine design, as well as in the overall understanding of immune mechanisms (Gomara and Haro, 2007). X-ray crystallography and Nuclear magnetic resonance techniques are trustable approaches for identifying BCEs (Mayer and Meyer, 2001). Nevertheless, these traditional experimental approaches are expensive and time-consuming (Kavitha, et al., 2013). The silico prediction tools can mitigate the identification workload by predicting epitope regions. For example, Jespersen et al. (Jespersen, et al., 2017) propose the commonly used tool BepiPred-2.0, which employs a random forest model to train annotated epitopes from antibody-antigen protein structures and then uses the trained model to predict newly generated antigen sequences. Afterward, with the increase in such biological data, a few deep learning methods are implemented for accurately predicting BCEs. EpiDope (Collatz, et al., 2021) uses a deep neural network to identify BCEs on individual protein sequences, which extras context-aware embeddings for each residue in the sequence by applying a vector with a length of 1000. In this way, EpiDope exceeds baseline methods in identifying BCEs. Although the above methods obtain good performance in identifying linear epitopes, they have difficulty in identifying conformational epitopes consisting of amino acid fragments that are far apart in the protein sequence but are brought together by the conformational folding of the polypeptide chain.

To solve these problems, several structure-based methods have been designed by considering spatial information. DiscoTope (Haste Andersen, et al., 2006) is the first method to focus explicitly on discontinuous epitopes by considering spatial information, amino acid statistics, and surface accessibility. DiscoTope-2.0 (Kringelum, et al., 2012) is the improved version of DiscoTope by summing propensity scores and half-sphere exposure as a surface measure. Nonetheless, DiscoTope-2.0 does not take into account glycosylation which may significantly affect epitopes. SEPPA 3.0 (Zhou, et al., 2019) investigates the impact of glycosylation in antigen surface patches, showing that antibodies may prefer to bind in N-glycosylation sites. ElliPro (Ponomarenko, et al., 2008) characterizes antigenic proteins by approximating them as ellipsoids and then calculates the protrusion index of the residues to cluster them. Epiope3D (da Silva, et al., 2022) is a novel scalable machine-learning method for accurately predicting conformational epitopes by using the concept of graph-based structural signatures. So far, structure-based tools have achieved decent performance, in general, better than sequence-based tools (da Silva, et al., 2022; Singh, et al., 2013). However, since experimentally decided structural information is usually not available, BCEs prediction must in many cases be conducted via amino-acid sequences alone.

With the great advances in deep learning technologies, protein structure prediction is undergoing a breakthrough. For example, AlphaFold2 (Jumper, et al., 2021) has integrated biological and physical knowledge about protein structure, information of multi-sequence alignments (MSA), and the complicated design of the deep learning model. AlphaFold2 has shown the ability to predict protein structure with atomic accuracy and proven accuracy competitive with experiments in a large number of cases in the 14th Critical Assessment of Protein Structure Prediction (CASP14). On the other hand, unsupervised pre-training using contextual language models has led to breakthrough improvements in natural language processing. Recently, these techniques have been employed in protein sequence representation learning and have shown very promising results in predictions including tertiary contacts, mutational effects, and secondary structure (Elnaggar, et al., 2020; Yuan, et al., 2022). These breakthroughs inspire us to develop an accurate B-cell epitope predictor by using the predicted protein structures and the pretrained language model.

In this study, we propose GraphBepi, a novel graph-based model for accurate epitope prediction. GraphBepi first generates the effective information sequence representations and protein structures from antigen sequences by the pretrained language model and AlphaFold2, respectively. GraphBepi then applies the edge enhanced deep graph neural network (EGNN) (Gong and Cheng, 2019) to capture the predicted protein structural information and leverages the bidirectional long short-term memory neural networks (BiLSTM) (Hochreiter and Schmidhuber, 1997) to capture long-range dependencies from sequences. The low-dimensional representation learned from EGNN and BiLSTM is then combined to predict B-cell epitopes. Through comprehensive tests on the curated epitope dataset, GraphBepi was shown to outperform the state-of-the-art methods.

## 2 Materials and methods

### 2.1 Dataset

To train and evaluate our model, we took a similar strategy as the study (da Silva, et al., 2022) to build a large epitope dataset. Specifically, we first fetched all biological assemblies with a resolution higher than or equal to 3 Å from Protein Data Bank deposited before 05/09/2022 (Berman, et al., 2000). Next, the ANARCI (Dunbar, et al., 2016) tool was used to identify antibody-antigen complexes and retain antigen chains with lengths of 25-1024. We labeled epitope residues in the antigen molecule depending on a cutoff distance standard, for example, any antigen residue having at least one heavy atom at a distance of less than 4 Å to an antibody residue will be treated as the epitope residue. We removed the antigen chain if it contained epitopes less than five. Next, we used MMseqs2 (Steinegger and Söding, 2017) to cluster antigen sequences, and any antigen sequence belonging to the same cluster was aligned to the cluster representative defined byMMseqs2 through tool blastp (Johnson, et al., 2008). Each clustering representative sequence was then modified as follows: If an epitope was labeled in the clustering representative, it would be kept. If an epitope was found on any antigen sequence of the aligned antigen sequences, it was transposed to the cluster representative sequence and marked as an epitope. This process was done at 95% sequence identity, resulting in the size of the dataset with 783 antigen sequences. We further conducted redundancy reduction via MMseqs2 at 70% sequence identity, resulting in generating a non-redundant data set of 633 antigen sequences. Finally, the antigen sequences deposited after 01/04/2021 were used as the independent test data (56 antigen sequences, consisting of 1393 binding residues and 14150 nonbinding residues, with 8.96% of epitope residues), and the rest of the antigen sequences were used as the training data (577 antigen sequences, consisting of 15981 binding residues and 119869 nonbinding residues, with 11.76% of epitope residues).

### 2.2 Protein representation

#### 2.2.1 Language model representation

The recent language model esm2_t36_3B_UR50D (Lin, et al., 2022) (denoted as ESM-2) is used for extracting features from each antigen sequence, which is an updated version of esm_msa1b_t12_100M_UR50S (denoted as ESM). The architecture of ESM and ESM-2 is based on the transformer model, and both of them are pretrained on UniRef50 (Suzek, et al., 2007) through the masked language modeling objective (Devlin, et al., 2018) in an unsupervised manner. We leverage the ESM-2 to extract sequence representation for each residue, which generates a 2560-dimensional feature vector for per-residue. We have also investigated another similar language model ProtT5-XL-U50 (Elnaggar, et al., 2020) (denoted as ProtTrans), which is first trained on BFD (Steinegger, et al., 2019) and then fine-tuned on UniRef50.

#### 2.2.2 Evolutionary information

Evolutionarily conserved residues probably include motifs correlated to crucial protein properties. For investigating the importance of evolutionary features, we test the widely used evolutionary features hidden Markov models (HMM) profile and position-specific scoring matrix (PSSM). The HMM profiles are produced by running HHblits (Remmert, et al., 2012) against UniClust30 (Mirdita, et al., 2017) using the default setting. The PSSM is generated by conducting PSI-BLAST (Altschul, et al., 1997) to seek the candidate sequence against UniRef90 (Suzek, et al., 2007) using an E-value of 0.001 and three iterations. Note that per-residue is embedded into a 20-dimensional vector through PSSM and HMM, respectively.

#### 2.2.3 Predicted protein structures

To take account of spatial information for each residue, we apply the predictive tool AlphaFold2 to predict protein structure. Specifically, we follow the tutorial at https://github.com/deepmind/alphafold to deploy AlphaFold2 on the Tianhe-2 supercomputer and then predict the protein structures. We have also investigated another similar protein structural prediction model, esmfold_v1 (Lin, et al., 2022) (denoted as ESMFold), which is a full end-to-end single sequence structure predictor. We download the pretrained ESMFold and then directly apply it to predict protein structures.

#### 2.2.4 Structural properties

We apply the program DSSP (Kabsch and Sander, 1983) to extract three types of structural properties from the AlphaFold2 predicted protein structures: (1) the profile of a nine-dimensional one-hot secondary structure, in which the first eight dimensions indicate the state of the eight-secondary structure, and the last dimension means the unknown secondary structure (2) Relative solvent accessibility (RSA) is obtained by normalizing the solvent accessible surface area (ASA) through the maximal possible ASA of the corresponding amino acid type. (3) Peptide backbone torsion angles PSI and PHI, which are transferred into a four-dimensional feature vector by cosine and sine transformations. In summary, these 13-dimensional structural feature vectors are named DSSP in this manuscript.

### 2.3 The architecture of GraphBepi

This study proposes a novel method GraphBepi for improving BCEs prediction by considering spatial information. As shown in Figure 1, the antigen sequence is fed into the pretrained language model and AlphaFold2 to generate the sequence embedding and protein structures, respectively. The relational graph of residues and DSSP are then extracted from protein structures. The sequence embedding and DSSP are then fed into the BiLSTM module to learn the effective representation by capturing long-range dependencies from sequences. They are also concatenated to form feature vectors of residues in the relational graph and are then fed into the edge-enhanced graph neural network (EGNN) to learn the structural information. Finally, the output of the EGNN and BiLSTM modules is concatenated to predict BCEs through a multilayer perceptron.

**Fig. 1.**
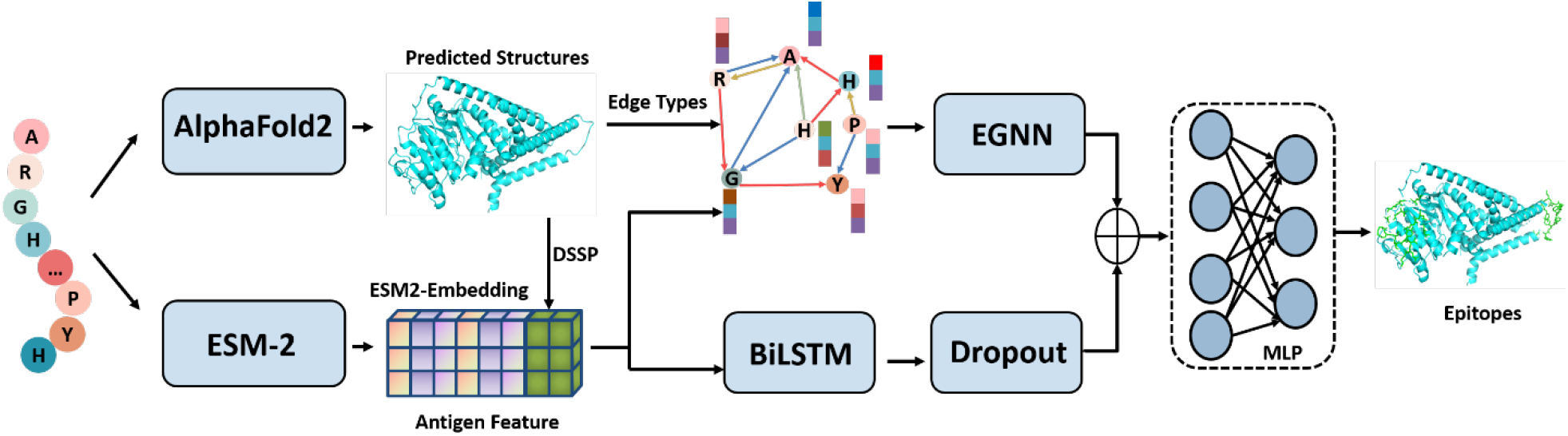
The network architecture of the proposed GraphBepi model. The input antigen sequence is respectively fed into the pretrained language model and AlphaFold2 to generate the ESM2-embedding and protein structures. The relational graph of residues and DSSP are extracted from the predicted protein structures. The ESM2-embedding and DSSP are then fed into the BiLSTM module to learn the effective representation by capturing long-range dependencies of the residues. They are also concatenated to form feature vectors of residues in the relational graph, which is then fed into the edge-enhanced graph neural network (EGNN) to learn the structural information. Finally, the output of the EGNN and BiLSTM modules is concatenated to predict BCEs through a multilayer perceptron.

#### 2.3.1 The Bidirectional LSTM module

The Long Short-Term Memory (LSTM) is a classic algorithm for capturing long-range dependencies, which is widely used in protein sequence encoding. The bidirectional LSTM (BiLSTM) model adds one more LSTM layer and reverses the direction of information. Briefly, it means that the input sequence flows backward in the additional LSTM layer. In this study, we use a bidirectional LSTM to process antigen sequences since they don’t have a specific direction. We denote the output of the BiLSTM as *H* for simplicity. The BiLSTM is employed for learning the DSSP structural properties obtained from predicted structures and the sequence embeddings obtained from ESM-2, respectively. We introduce the BiLSTM algorithm carefully in Supplementary Note 1.

#### 2.3.2 Graph construction

We build the residue-level relational graph *G* = (*N, ε, R*) for the structure of an antigen following the ref (Zhang, et al., 2022), where *N* and *ε* mean the set of nodes and edges respectively, and *R* is the set of edge types. The term (*i, j, r*) is used for indicating the edge from node *i* to node j with edge type r. We add three types of directed edges into the graph including K-nearest neighbor edges, radius edges, and sequential edges. They are generated as follows: (1) Sequential edges: residues *i* and *j* will be connected by an edge if |*j* – *i*| < *d_seq_*, where |*j* – *i*| represent the sequential distance between residues *i* and *j*, and *d_seq_* is a predefined threshold. Then, the *d* = *j* – *i* is used as the edge type between residue *i* and *j*. Hence, there are 2*d_seq_*-1 types of sequential edges. (2) Radius edges: the radius edges between two residues are also added if the spatial distance between them is less than a threshold *d_radius_*. (3) K-nearest neighbor edges: KNN edge types are added by computing the k-nearest neighbors based on the Euclidean distance. In this study, the sequential distance threshold *d_seq_* and the radius are set to 3 and 10, respectively. The number of residue neighbor k is set to 10. Finally, the edge types consist of radius edges, KNN edges, and sequential edges, resulting in *2d_seq_*+1=7 different types of edges.

#### 2.3.3 Edge enhanced graph neural network (EGNN) module

We apply the EGNN (Gong and Cheng, 2019) framework to capture spatial information by adequately integrating node (residue) features and multiple-dimensional edge features. Given a residue graph with *N* residues, we first let *X* be an *N* × *F* matrix representing the node features of the entire graph, where *F* is the dimension of the node feature. The edge features are represented by an *N* × *N* × *P* tensor, where *P* is the dimension of the edge feature. Therefore, E_*ij*_-indicates the P-dimensional feature vector of the edge that connects residue *i* and residue *j*. E_*ijp*_ represents the *p*-th channel of the edge feature in E_*ij*_. Concretely, the feature vector 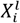 of residue *i* at the *l* layer will be aggregated from feature vectors of the neighboring nodes by simultaneously integrating the edge features. The aggregation operation is represented as follows:

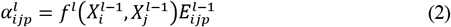

where *σ* is the activation function. *α* is the attention operation, which is guided by edge features of the edge connecting two residues; *α_..p_* represents the *p* channel matrix slice of the *N* × *N* × *P* tensor. Specifically, we treat multiple-dimensional edge features as multi-channel signals and then conduct a separate attention operation for each channel. These results from each channel are then combined by concatenation operation *||*. For an individual channel of edge features, the attention function can be done as follows:

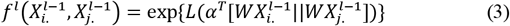

where *f^l^* is the attention function implemented by a linear function for simplicity:

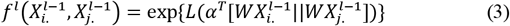

where *L* is the LeakyReLU activation function. || is the concatenation operation. In Equation (1), *g* is the transformation function mapping the node features from the input space to the output space, which can be formulated as follows:

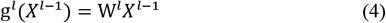

where W^*l*^ is an *F^l^* × *F*^*l*−1^ parameter matrix. Finally, the attention coefficients will be treated as the new edge features for the next layer. By doing so, EGNN efficiently integrates the edge features and node features.

#### 2.3.4 Multilayer perceptron

We combine the output of the EGNN module and the BiLSTM module via concatenation operation, and then feed them into the multilayer perceptron (MLP) to predict the B-cell epitope binding probabilities of all *N* amino acid residues as follows:

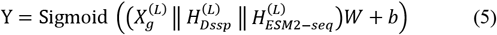

where 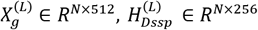 and 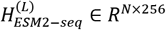 are the output of the last layer of EGNN module and the BiLSTM module, respectively. *Y* ∈ *R*^*N*×1^ is the predictive result of *N* amino acid residues.

### 2.4 Implementation details

We conducted 10-fold cross-validation (CV) for the training dataset, where the dataset was split into 10-folds randomly. Each time, our method was trained on 9-folds and then evaluated on the one rest fold. This process was performed ten times. The performance of our method on the 10-folds was averaged as the final validation performance. For the testing dataset, all ten trained models in the CV were used for making predictions, which were then averaged as the final testing predictions. In addition, the number of layers and hidden units of BiLSTM were set to 3 and 128, respectively. For the EGNN module, the number of layers and hidden dimensions were set to 2 and 256, respectively. We leveraged the Adam optimizer with a batch size of 4 and a learning rate of 1e-6 for model optimization on the binary cross-entropy loss. Our method was implemented through Pytorch and python. We fixed the training epoch to 300 epochs since the average performance of the validated data for the 10 trained models was the highest at this epoch. All experimental results were conducted on an Nvidia GeForce RTX 3090 GPU.

### 2.5 Evaluation metrics

Similar to the previous study (Yuan, et al., 2021), six commonly used metrics are employed for evaluating prediction performance. They are precision (Pre), recall (Rec), F1-score (F1), Matthews correlation coefficient (MCC), area under the precision-recall curve (AUPR), and area under the receiver operating characteristic curve (AUC):

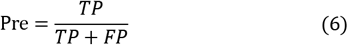

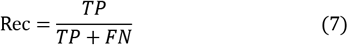

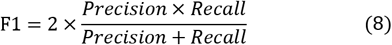

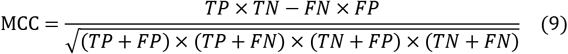

where true negatives (TN) and true positives (TP) represent the number of non-binding and binding residues predicted correctly, respectively. False negatives (FN) and false positives (FP) represent the number of incorrectly identified non-binding and binding residues, respectively. AUPR and AUR are independent of thresholds, therefore revealing the overall performance of the model. The rest metrics are computed through a predefined threshold to convert the predicted binding probabilities to binary predictions, where the threshold is decided by maximizing the F1 score.

## 3 Results

### 3.1 Performance on the 10-fold CV and independent test

To evaluate the performance of GraphBepi, we tested it by AUC, AUPR, F1, and MCC using the 10-fold CV and independent test data. Concretely, the GraphBepi model achieved AUC values of 0.723 and 0.751, AUPR values of 0.245 and 0.261, F1 values of 0.320 and 0.310, and MCC values of 0.212, and 0.232 on the 10-fold CV and the independent test, respectively (Supplementary Table S1). The consistent performance on the CV and independent test demonstrated the robustness of our model. For further investigating the advantages of antigen geometric information and the edge enhanced graph neural network, we compared GraphBepi with a baseline model transformer consisting of two-layer networks. The transformer was used as a geometric agnostic baseline to test the impact of the structural information for binding residue prediction, which was fed the same input features as GraphBepi. As shown in Supplementary Table S2, GraphBepi consistently outperformed than the baseline model transformer, which were 4.6%, 6.4%, 5.1%, and 5.6% higher than transformer in terms of AUC, AUPR, and F1, MCC. Here, the edge enhanced graph neural network helped GraphBepi focus on the spatially adjacent residues, which learned the remote residues connected in the whole graph by efficiently integrating different edge features and residue features. As shown in Supplementary Table S2, the removal of edge features caused a decrease of 2.9% in terms of AUPR on the independent test.

The improved performance of our method over the transformer may be due to its better ability in capturing spatial information. To further investigate, we evaluated the performance of our method and transformer on amino acids with different numbers of non-local contacts. We considered two residues as non-local contacts if the distance between these two residues was greater than 20 residues in sequence position and less than 12 Å in their Cα atomic distance. Figure 2 showed that our method consistently outperformed the transformer on the independent test data. Concretely, the performance of our method outperformed the transformer by 11% in terms of F1 on the amino acids with 0-9 non-local contacts, and the performance gap expanded to 52 % when the number of non-local contacts were larger than 20. Similar trends could be found when measured by other metrics (Supplementary Figure S1). The results demonstrated that GraphBepi efficiently captured the spatial information of antigen structures.

**Fig. 2.**
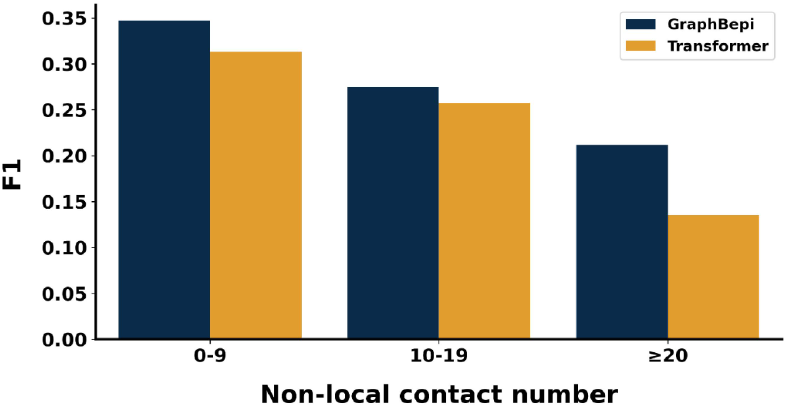
The F1 values of GraphBepi and transformer on amino acids with the different numbers of non-local contacts.

### 3.2 Representation from pretrained language models is informative for predicting BCEs

To evaluate the effect of sequence representation extracted by the ESM-2 language model used in the GraphBepi, we compared ESM-2 with the widely used handcrafted features in this field, and another two language models ESM and ProtTrans. As shown in Table 1, on the independent test, our model gained an average AUC and AUPR of 0.736 and 0.240 respectively when only using the sequence representation extracted from ESM-2. Whereas, the evolutionary information (HMM and PSSM, EVO) and the DSSP led to a decrease in the performance of our method (4.9% and 6.9% decrease in terms of AUPR for EVO and DSSP, respectively). The results demonstrated that the embeddings learned from the pretrained language model ESM-2 were better than the handcrafted features.

**Table 1.** The predictive performance on the independent test using different features.

| Feature group | AUC | AUPR | F1 | MCC |
| --- | --- | --- | --- | --- |
| EVO | 0.717 | 0.191 | 0.269 | 0.185 |
| DSSP | 0.680 | 0.171 | 0.235 | 0.142 |
| ESM-2 | 0.736 | 0.240 | 0.291 | 0.212 |
| EVO + DSSP | 0.723 | 0.200 | 0.271 | 0.192 |
| ProtTrans + DSSP | 0.750 | 0.260 | 0.309 | 0.214 |
| ESM + DSSP | 0.744 | 0.216 | 0.293 | 0.212 |
| ESM-2 + EVO + DSSP | 0.757 | 0.254 | 0.317 | 0.240 |
| ESM-2 + DSSP(GraphBepi) | 0.751 | 0.261 | 0.310 | 0.232 |

Note: EVO means the evolutionary features HMM and PSSM.

In addition, we further investigated the performance of our model when integrating different features. We noted that the performance of EVO could be improved by integrating the structural properties DSSP. Therefore, we also integrated DSSP with ESM-2, which improved by 8.75% in terms of the AUPR metric. These results demonstrated that the structural properties of amino acids such as secondary structure and RSA were sufficient to reserve the complicated patterns of epitopes. We then integrated more information including evolutionary features, ESM-2 embeddings, and DSSP. The results showed that combing all information brought a minor improvement in the test data (< G.0.01 of AUC). These results demonstrated that the ESM-2 language model may potentially reserve the evolutionary information of the protein. We found ESM-2 performed slightly better than ProtTrans, which may be because they took different network architectures. ESM-2 and ProtTrans outperformed ESM in all metrics, probably because they used more training parameters and data sets.

### 3.3 Comparison with state-of-the-art methods

In this section, we investigated the relative importance of each module in GraphBepi (Table 2) and compared our method with state-of-the-art methods (Table 3). We first compared GraphBepi with two sequence-based methods (Bepipred 2.0 and EpiDope) and four structure-based approaches (ElliPro, Discotope 2.0, epitope3D, and ScanNet (Tubiana, et al., 2022)) using their default parameters. ScanNet provided the option to select a transfer learning strategy or not. In this study, ScanNet was called Scan-Net_T when a transfer learning strategy was used, otherwise it was called ScanNet_WT. As shown in Table 3, our method surpassed the second-ranked method ScanNet_T by 5.5% in terms of AUC, 44.0% in terms of AUPR, 20.4% in terms of F1, and, 36.9% in terms of MCC, respectively. We noted that ScanNet_T achieved higher performance than Scan-Net_WT, probably because the transfer learning strategy generated profitable parameters for the initial model. Discotope-2.0 was the third-ranked method, which achieved comparable performance with method ElliPro in terms of AUC value. The AUPR value of Discotope-2.0 was 3.2% higher than that of ElliPro. However, both of them performed lower than Scan-Net_WT, which may be because ScanNet_WT applied the deep learning network architecture and transfer learning strategy. Interestingly, although epitope3D was a structure-based method, it performed lower than the sequence-based method EpiDope. The reason may be that the experimental structures also brought noises due to the flexible characteristic of protein structure. Note that GraphBepi did not have the highest recall since it was an unbalanced measure strongly relying on thresholds.

**Table 2.** The predictive performance of GraphBepi on the independent test when removing each module.

| module | AUC | AUPR | F1 | MCC |
| --- | --- | --- | --- | --- |
| GraphBepi w/o EGNN | 0.706 | 0.209 | 0.275 | 0.194 |
| GraphBepi w/o BiLSTM | 0.742 | 0.239 | 0.300 | 0.226 |
| GraphBepi | 0.751 | 0.261 | 0.310 | 0.232 |

Note: w/o represents without the corresponding module

**Table 3.** Performance comparison of GraphBepi with state-of-the-art methods on the independent test data.

| Method | AUC | AUPR | F1 | MCC | Rec | Pre |
| --- | --- | --- | --- | --- | --- | --- |
| EpiDope | 0.547 | 0.102 | 0.173 | 0.046 | 0.855 | 0.096 |
| Bepipred-2.0 | 0.648 | 0.132 | 0.220 | 0.126 | 0.562 | 0.137 |
| ElliPro | 0.632 | 0.122 | 0.217 | 0.123 | 0.592 | 0.133 |
| Discotope-2.0 | 0.655 | 0.154 | 0.231 | 0.136 | 0.405 | 0.162 |
| epitope3D | - | - | 0.135 | 0.039 | 0.176 | 0.109 |
| ScanNet_WT | 0.648 | 0.135 | 0.218 | 0.121 | 0.532 | 0.137 |
| ScanNet_T | 0.712 | 0.182 | 0.257 | 0.170 | 0.440 | 0.182 |
| GraphBepi | 0.751 | 0.261 | 0.310 | 0.232 | 0.393 | 0.255 |

Note: ScanNet\_T means that method ScanNet uses the transfer learning strategy, while ScanNet\_WT means that it does not.

To further investigate the advantages of our method, we analyzed the relative importance of each module in GraphBepi by conducting the model ablation study on the test data. As shown in Table 2, the removal of the BiLSTM module caused a decrease of 0.9% and 2.2% in terms of AUC and AUPR. This change indicated that the BiLSTM could capture long-range dependencies of amino acid residues. The removal of the EGNN caused the greatest decrease of 4.5% and 5.2% in terms of AUC and AUPR. The results demonstrated that the spatial information was efficiently captured by the EGNN module. In summary, the cooperation of each module achieved the best performance.

### 3.4 Impact of the quality of predicted protein structure

Since our method applied predicted protein structures, the quality of protein structures predicted from structural prediction models might be essential to the downstream BCEs prediction. Here, we first evaluated the performance of our model by using the native protein structures and the predicted protein structures from AlphaFold2 and ESMFold. As shown in Table 4, similar results were obtained when our method used protein structures predicted from AlphaFold2 and native structures. Concretely, the AUC and AUPR of native structures were only 0.8% and 0.2% higher than the predicted structures from AlphaFold2. In addition, the performance of protein structures predicted from AlphaFold2 was higher than the protein structures predicted from ESMFold, which might be ascribed to the different architectures they applied for predicting protein structures. On the other hand, our method achieved the lowest performance when only using the sequence protein, which demonstrated that the structural information was important for BCEs prediction.

**Table 4.**
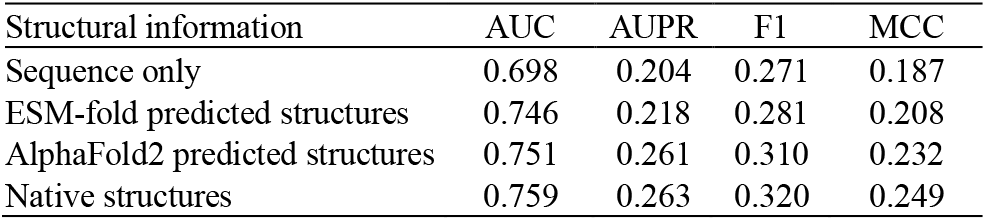
The predictive performance of GraphBepi when using only sequence, or different predicted structures.

| Structural information | AUC | AUPR | F1 | MCC |
| --- | --- | --- | --- | --- |
| Sequence only | 0.698 | 0.204 | 0.271 | 0.187 |
| ESM-fold predicted structures | 0.746 | 0.218 | 0.281 | 0.208 |
| AlphaFold2 predicted structures | 0.751 | 0.261 | 0.310 | 0.232 |
| Native structures | 0.759 | 0.263 | 0.320 | 0.249 |

To further investigate the advantages of geometric deep learning employed in our method by using the predicted protein structures from AlphaFold2, we computed the average global distance test (Zemla, 2003) (called GDT) between the native structures and the predicted structures through the tool Spalign (Yang, et al., 2012). Figure 3 showed the quality of the predicted protein structures and the F1 values of each antigen on the independent test data for GraphBepi (black scatters). Specifically, we sorted the antigens based on GDT value and then divided them into nine bins evenly to calculate the mean GDT and F1 for every bin (red line). The results showed that positive correlation between the AlphaFold2 predicted quality measured by GDT and the GraphBepi performance measured by F1 on an independent test set. The top 20% of antigens with the highest GDT (mean GDT=0.974) had a mean F1 of 0.406. Whereas, the bottom 20% of antigens with the lowest GDT (mean GDT=0.563) had a mean average F1 of 0.241, which was lower than that of the top 20% of proteins. Similar trends could be found in terms of other metrics as shown in Supplementary Figure S2. These results suggested the importance of the quality of antigens structures predicted by AlphaFold2 for BCEs prediction.

**Fig. 3.**
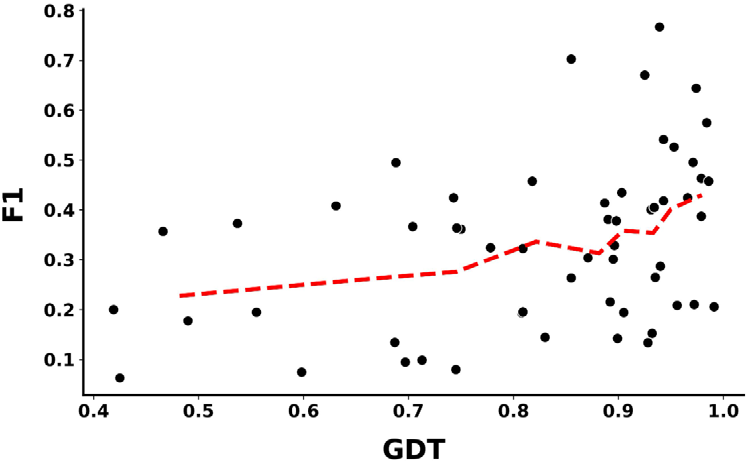
Positive correlation between the AlphaFold2 predicted quality measured by GDT and the GraphBepi performance measured by F1 on the independent test set. The black scatter indicates the GDT and F1 for each protein, while the red line indicates the average GDT and F1 for each bin after sorting all antigens by GDT and dividing them into nine bins.

### 3.5 Case study

To demonstrate the superiority of GraphBepi, we visualized one example (PDB ID: 7S2R, chain A) from the independent test for illustration. Figure 4 showed the BCEs predictive results of GraphBepi (a) GraphBepi without EGNN (b), the second-ranked method ScanNet_T (c), and the sequenced-based method Bepipred-2.0 (d). This case antigen sequence included 17 epitopes over a total of 197 residues. GraphBepi predicted 37 binding residues, of which 13 were TP, resulting in an AUPR of 0.558, F1 of 0.481, and MCC of 0.454. By comparison, the GraphBepi without EGNN predicted 93 binding residues, of which 16 were TP, resulting in a lower AUPR of 0.451, F1 of 0.291, and MCC of 0.289. The results showed that the spatial information captured by the EGNN module could help our method accurately identify the epitopes and reduce the false positive rate. A similar trend could be found in ScanNet_T and Bepipred-2.0. Concretely, the structure-based method ScanNet_T predicted 57 binding residues, of which 11 were TP, resulting in an AUPR of 0.332, F1 of 0.297, and MCC of 0.242. Whereas the sequenced-based Bepipred-2.0 predicted 87 binding residues, of which 12 were TP, resulting in a lower AUPR of 0.127, F1 of 0.226, and MCC of 0.157. In addition, the visualization results for other competing methods can be found in Supplementary Figure S3, and they also performed worse than our method.

**Fig. 4.**
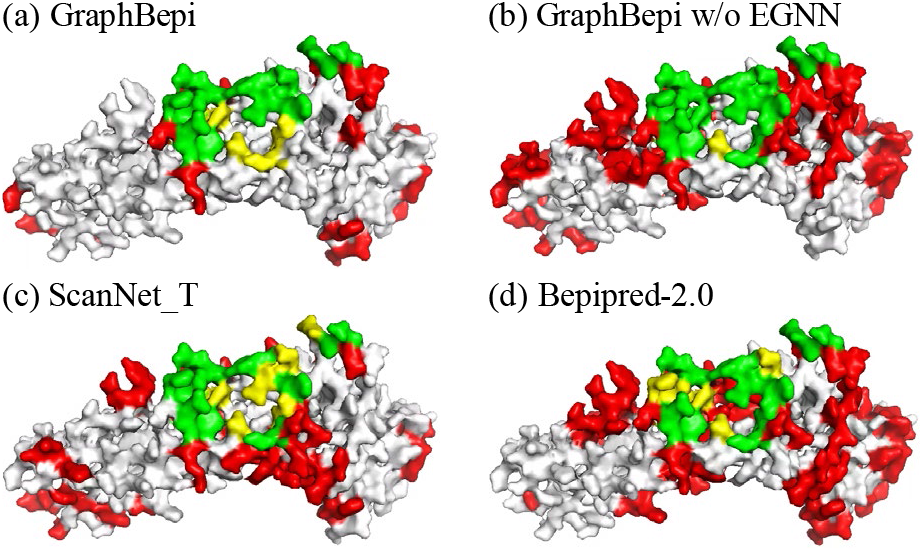
Visualization of an example (PDB ID: 7S2R, chain A) from the test data predicted by GraphBepi (a), GraphBepi without EGNN (b), ScanNet_T (c), and Bepipred-2.0 (d). True positives, false negatives, and false positives are colored in green, yellow, and red, respectively.

## 4 Discussion

Identifying the B-cell epitopes is an essential step for guiding rational vaccine development and immunotherapies. Here, we propose GraphBepi, a novel graph-based model for accurate epitope prediction by using structural information predicted from AlphaFold2. GraphBepi applies the edge-enhanced deep graph neural network (EGNN) to capture the predicted protein structures and leverages the bidirectional long short-term memory neural networks (BiLSTM) to capture long-range dependencies from sequences. The low-dimensional representation learned from EGNN and BiLSTM is then combined to predict B-cell epitopes. Through comprehensive tests on the curated epitope data set, GraphBepi was shown to outperform the state-of-the-art methods.

In spite of several sequence-based methods also are designed for identifying the BCEs such as EpiDope and Bepipred 2.0. Whereas they achieve limited performance since they only use the contextual features of the sequential neighbors. Structure-based methods try to solve these problems by considering spatial information. However, they are not applicable to most proteins that don’t have known tertiary structures. More importantly, these methods only improve the predictive performance by using hand-crafted evolutionary information. Therefore, we develop the method GraphBepi for predicting BCEs by leveraging spatial information predicted from AlphaFold2 and using the effective embeddings extracted from the pretrained language model. Benefitting from the dedicated design, our method achieves the best performance compared to the state-of-the-art methods. On the other hand, our method achieves similar inference times (about 0.1 seconds for one antigen) to the best competing method.

Despite the advantages of GraphBepi, our method can be improved in several aspects. For example, the performance of our approach is influenced by the quality of antigen structures predicted from AlphaFold2. This might be relieved by including other informative sequence-derived features to increase the robustness of the model. We will explore these challenges in future work. In summary, we demonstrate that GraphBepi provides a deep-learning model for accurately predicting BCEs. GraphBepi is also freely available in an easy-to-use web interface at https://bio-med.nscc-gz.cn/apps/GraphBepi.

## Funding

This study has been supported by the National Key R&D Program of China (2020YFB0204803), National Natural Science Foundation of China (61772566 and 62041209), Guangdong Key Field R&D Plan (2019B020228001 and 2018B010109006), Introducing Innovative and Entrepreneurial Teams (2016ZT06D211) and Guangzhou S& Research Plan (202007030010).

## Notes

### Competing Interest Statement

The authors have declared no competing interest.

